# The *Hoja loca1* Mutant and AUX/IAA Function in Phytomer Development

**DOI:** 10.1101/2020.03.27.012211

**Authors:** A. E. Richardson, A. Sluis, S. Hake

## Abstract

Plant architecture is determined by the iterative building of phytomer units, each containing a portion of stem, an organ and an axillary meristem. Each phytomer can follow different developmental paths underpinning the complexity, and plasticity, of plant form. Auxin plays a central role in the coordination of phytomer development, regulating organ initiation and patterning across all axes. This diversity in auxin function results from changes in the activities and expression of auxin signaling components, including the AUX/IAA repressors. Higher land plants have multigene AUX/IAA families, which leads to functional redundancy. Dominant mutations, which prevent AUX/IAA degradation in response to auxin, have highlighted the importance of these proteins in fine-tuning organ development. Here we report a new dominant AUX/IAA mutant in maize, *Hoja loca1* (*Oja*). *Oja* has a mutation in the degron motif of ZmIAA28 and specifically affects aerial organ initiation and leaf medio-lateral patterning, but phytomer initiation remains unchanged. Mutant phenotypes are variable and transcriptional profiling of individual meristems identified clusters of genes that may underpin the phenotypic differences. The unique phenotype of *Oja* provides evidence of species-specific sub-functionalization of the AUX/IAAs, and illustrates the crucial role of auxin signaling in the tight coordination of phytomer unit development.

## Introduction

The iterative post-embryonic development of plants has led to impressive developmental plasticity, which allows plants to modify their architecture over time and respond to environmental conditions. Organs initiate from meristems, groups of self-organizing totipotent cells, in regular phyllotactic patterns. Initiation of each leaf includes an axillary meristem and an internode. This developmental unit of leaf, axillary meristem, and internode is known as a phytomer. Aerial plant development can therefore be understood as the successive formation of phytomer units (Galinat, 1959), with developmental plasticity manifesting in different amounts of growth in the three components.

The fate of cells within the meristem is dependent on position. Cells in the center of shoot meristems divide slowly, are isodiametric in shape, and are considered indeterminate. As divisions push cells from the center outward, cells enter the peripheral or morphogenetic zone, where organs initiate. It is here that the primordium is specified, forming a leaf at plastochron 0 (P_0_) (Bolduc et al., 2012a). The *KNOTTED1-LIKE HOMEOBOX* (*KNOX*) genes mark indeterminate cell fates (Jackson et al., 1994; Smith et al., 1992), and are down-regulated in P_0_ cells. *KNOX* gene expression is maintained within the other components of the phytomer, the internode and bud, until they elongate (Tsuda et al., 2017). The downregulation of *KNOX* genes in the P_0_ is preceded by the formation of an auxin maximum, coordinated by a convergence point of the auxin transporter PINFORMED1 (PIN1) in the epidermis (Bayer et al., 2009; Benková et al., 2003; O’Connor et al., 2014; Reinhardt et al., 2003). Auxin acts by triggering the degradation of AUX/IAA proteins, allowing AUXIN RESPONSE FACTORS (ARFs) to regulate downstream genes (Lavy and Estelle, 2016; Salehin et al., 2015). These coordinated events comprise the mechanism for organ initiation.

As organs develop and differentiate, asymmetries are established that determine tissue identity across proximal/distal, abaxial/adaxial, and medial/lateral axes (Kuhlemeier and Timmermans, 2016; Richardson and Hake, 2019). In grasses, the leaf initiates as a ring-shaped primordium around the meristem with the separation of the lower leaf (sheath) margins occurring later in development (Johnston et al., 2015; Sharman, 1942). Proximal-distal patterning in the grass leaf leads to the formation of a distal blade and a proximal sheath. The ligule and auricle form at the boundary of blade and sheath. For most grasses, the medial/lateral axes are marked by a midvein at the center which is thickened in comparison with the margins of the leaf (referred to as the midrib).

Auxin signaling has been implicated in all stages and axes of grass leaf development. Treatment of maize shoot apices, with the auxin inhibitor NPA (*N-1*-Naphthylphthalamic Acid) limits organ initiation and prevents the separation of the sheath margins resulting in a tube leaf (Scanlon, 2003). PIN proteins are also expressed at the position of a ligule prior to its elaboration (Moon et al., 2013) and RNAseq analysis of leaf microdomains has shown that ligule development involves the recapitulation of the organ initiation mechanism (Johnston et al., 2014). The establishment of the abaxial/adaxial axis also involves auxin signaling, utilizing opposing gradients of tasiR-ARFs and miR166 (Nogueira et al., 2007). Despite the role of auxin signaling throughout grass leaf development, auxin signaling mutants (i.e. mutants in AUX/IAA or ARFs), which affect leaf development, have not yet been reported in grasses.

Here we describe a dominant mutant, named *Hoja loca1* (*Oja*), with a range of leaf defects. In its most severe expression, mutants fail to initiate leaves, although an internode will be partitioned to each phytomer. In the least severe, midveins are missing from otherwise correctly patterned leaves. Between these extremes, tube leaves are formed, where the leaf margin is completely fused. Remarkably, leaves maintain proximal distal and abaxial/adaxial patterning. Using positional cloning and whole genome sequencing, we identify the gene as the AUX/IAA protein, ZmIAA28 (GRMZM2G035465). By analysing individual meristems, we were able to use the variable phytomer phenotypes of *Oja* to dissect potential regulatory roles of ZmIAA28 that may be responsive to different auxin concentrations. Unlike other maize AUX/IAA mutants that specifically affect roots (Woll et al., 2005) or inflorescences (Galli et al., 2015), *Oja* provides a view into the unique role of AUX/IAA proteins in grass leaf patterning. The ability of *Oja* to produce a leafless phytomer also hints at the role of auxin signaling in coordinating development of the separate phytomer components.

## Results

### Hoja loca1

*Hoja loca1* (*Oja*) is a dominant mutant that was discovered in a forward genetics EMS screen. The phenotype is variable, ranging from mild to severe, and it exhibits unpredictable expression through generations; crosses made with mildly affected plants give rise to severe plants and vice versa.

### *Hoja loca1* leaves have medial/lateral defects but normal proximal/distal and abaxial/adaxial patterning in aerial organs

Maize leaves are divided into a proximal sheath that wraps around the inner leaves and a distal blade that bends away from the plant, optimized for photosynthesis (Figure 1Ai). At the junction of the blade and sheath is the ligule, an epidermal fringe on the adaxial surface and two auricles along the margin that allow the blade to tilt away from the sheath (Figure 1Aii-iv). The blade contains a midrib, a thickened region with chlorophyll-containing cells on the abaxial surface and clear cells on the adaxial surface (Figure 1Av), that supports the extended blade.

**Figure 1.**
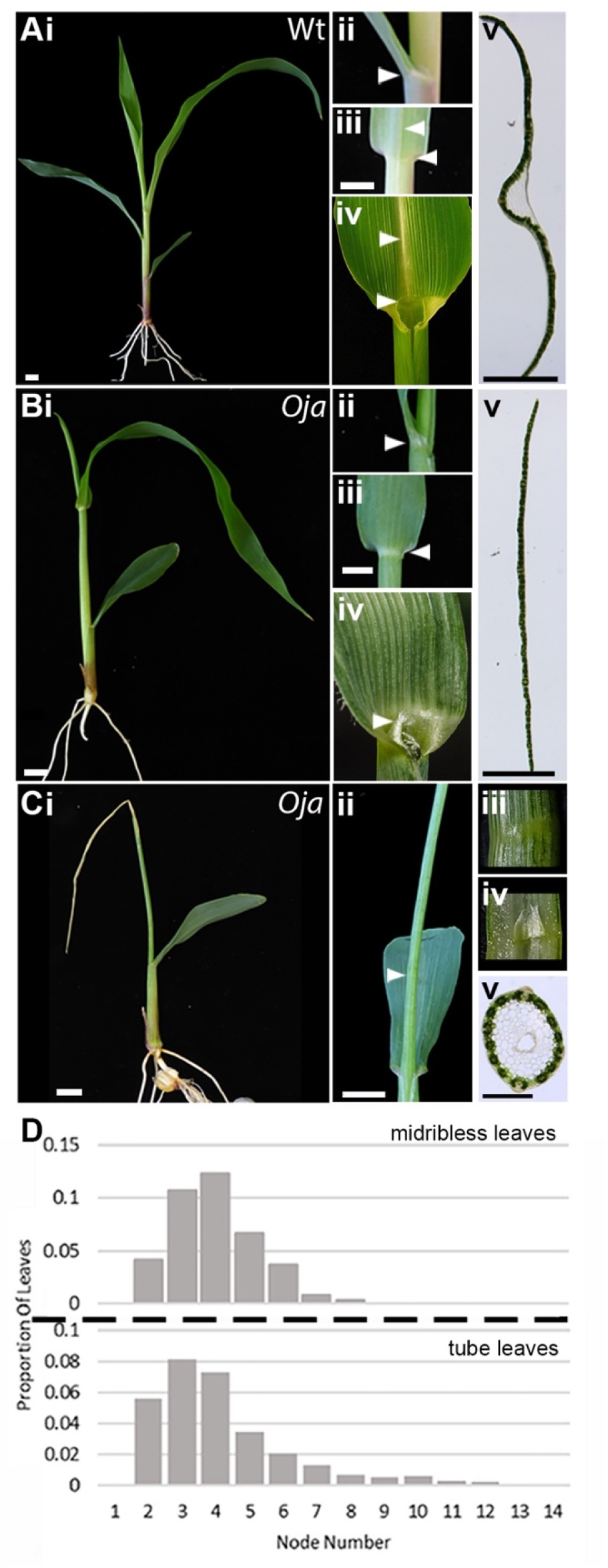
Hoja loca1 mutants have leaf defects. The phenotype of wild type (Wt) (A), mild (B) and severe (C) *Oja* mutants including (i) whole-seedling, and (ii-iv) magnified images of the ligule/ auricle regions, showing the side (ii) and abaxial (iii) and the adaxial (iv) views. (C iv) is a cross-section through the tube leaf showing the intact ligule (arrowhead in Cii). (v) Cross-sections through seedling leaves 1cm above the ligule. Scale bars are 1cm, arrowheads indicate the ligule or midrib regions. (D) The distribution of midribless and tube leaves in *Oja* plants (n=150), node 1 is at the base of the plant.

In contrast to wild-type seedlings, *Hoja loca1* (*Oja*) mutants have varied leaf phenotypes present on the same plant, including normal leaves (Figure 1B,C). The mildest leaf mutant phenotype has missing midribs (leaf 2 in Figure 1Bi). These leaves droop backwards without the structural support of the midrib (Figure 1Bi-v). The midribless leaves completely lack adaxial clear cells, although the flanking blade regions appear normal (Figure 1Bv). Midribless leaves still have normal ligules and auricles (Figure 1Bii-iv, arrowheads) suggesting that the midrib is not essential in elaboration of proximal/distal patterning.

Severe mutant leaf phenotypes include tube leaves in which the margins are not separated (Figure 1C). As in the case for midribless leaves, tube leaves also have normal proximal/distal patterning, developing a normal ligule between the sheath and blade (Figure Cii-iv). Cells in the tube leaves also have correct abaxial/adaxial polarity with chlorophyll containing cells on the abaxial surface and clear cells on the adaxial surface (Figure 1Cv). These results suggest that *Oja* does not influence proximal/distal or abaxial/adaxial patterning.

To determine whether there was a pattern in initiation of different leaf phenotypes, we recorded the occurrence of midribless and tube leaves in a population of 150 plants (Figure 1D). Both midribless and tube leaves can be observed in the same plant. In all cases, the first leaf is never affected. The second leaf may show one of these defects and the third and fourth are most likely to be mutant (Figure 1D). The frequency for both defects decreases during development but tube leaves could be observed late in development. This variability of phytomer phenotype makes *Oja* difficult to work with, but also provides a unique opportunity to dissect the different functions of the underlying gene.

In many *Oja* plants, inflorescence branching and floral development was severely affected. Both male and female inflorescences bore bare regions indicative of a failure to initiate spikelet primordia, and the tassels were not branched. However, when flowers did form, they appeared to be normal and fertile (Supplementary Figure 1). In contrast to aerial development, root development in seedlings appeared to be normal with initiation of primary, seminal and lateral roots and a normal gravitropic response (Supplementary Figure 2). *Oja* therefore specifically affects aerial organ development.

### *Hoja loca1* leaves have defects in vein distribution, but normal vein density

Given the midrib defect, we next questioned whether veins are altered in *Oja* mutants. Seedling leaf cross-sections showed that the abaxial/adaxial patterning of vascular bundles appears normal (Figure 2A,B), mirroring the normal abaxial/adaxial patterning of the entire organ.

**Figure 2.**
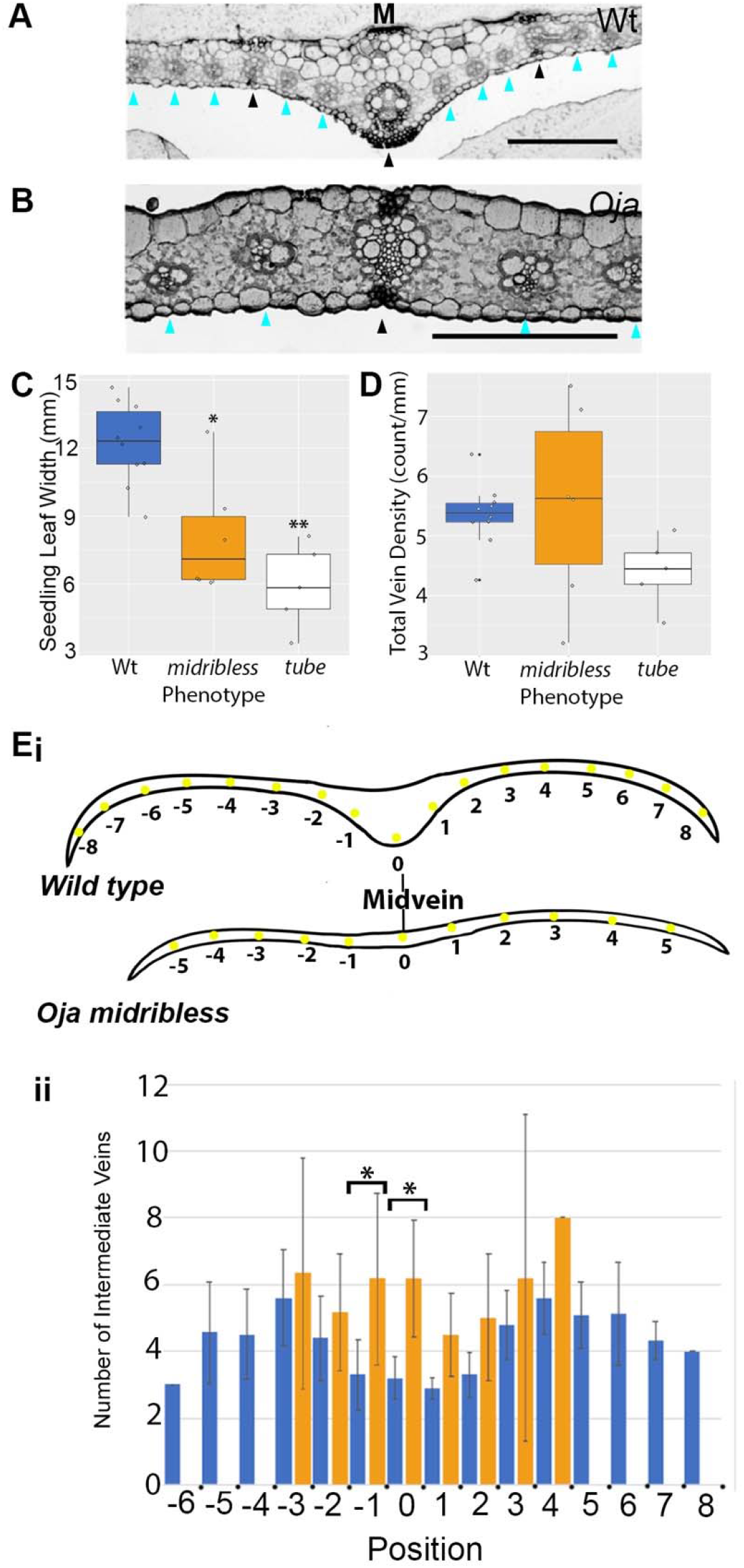
Hoja loca1 mutants have a defect in vein distribution. (A-B) Plastic sections through wild type (A) and midribless (B) leaves. The midrib is indicated by the M, the lateral veins by the black arrowheads, and intermediate veins by the cyan arrowheads. Scale bars are 1mm. (C) Seedling leaf width varies significantly between wild type and midribless (n=10 of each phenotype, ANOVA and Tukeys HSD, df=2, * p= 0.0032) and between wild type and tube leaves (**, p=0.000083). (D) Total leaf vein density does not vary significantly. (E) Leaf vein distribution is perturbed in midribless mutants. (Ei) Cartoons depicting the alignment of midribless and wild-type leaves. Wild type and mutant leaves were aligned by allocating the position of “0” to the wild type midvein, and the lateral vein closest to the midpoint of the leaf in the midribless mutant. Each successive lateral vein was then allocated a position relative to 0 (Ei) and the number of intermediate veins between each position was recorded. (Eii) The mean number of intermediate veins near the midvein were significantly different in the midribless mutant (orange) when compared to wild type (blue) (n=10 for each phenotype, ANOVA and Tukeys HSD, * p = 0.0051 and p=0.00089).

To assess whether the medio-lateral patterning of vascular bundles was altered we measured leaf width and determined vein density and distribution in the third leaf of 3-week-old seedlings. Midribless leaves were narrower than wild type and tube leaves were less than half the width of wild type (Figure 2C). Overall vein density was not affected by *Oja* (Figure 2D) suggesting a reduction in total vein number to match the decrease in size. To assess vein distribution, we counted the number of intermediate veins between lateral veins in both wild-type and midribless leaves. In wild type, the number of intermediate veins shows a symmetry across the leaf, with a dip at the mid position (Figure 2Eii, blue). We did not identify a dip of intermediate veins in the midvein position in *Oja* midribless leaves (Figure 2Eii, orange), supporting the hypothesis that medio-lateral patterning is affected in the *Oja* mutant.

### *Hoja loca1* mutants fail to initiate leaves but still set aside internode cells

At each leaf initiation, cells are set aside for the leaf and the internode below (Johri and Coe, 1983). A mature maize plant is thus composed of alternating internodes and nodes, which is the position of leaf insertion. In mature *Oja* plants we observed bare nodes, for example the plant in Figure 3A ended in a tube leaf after four nodes failed to elaborate leaves (arrows). To assess whether these bare nodes were due to ectopic nodes or a failure of leaf initiation, we counted the number of nodes in *Oja* versus wild-type siblings and compared these to the number of leaves. In mature plants there is no significant difference in number of nodes (Figure 3B), but there are significantly fewer leaves in *Oja* mutants (Figure 3C). This alternating node, internode, node pattern without accompanying leaves is also observed when slicing through developing tube leaves (Figure 3D). These data suggest that *Oja* may be able to initiate new phytomers consisting of just nodes and internodes.

**Figure 3.**
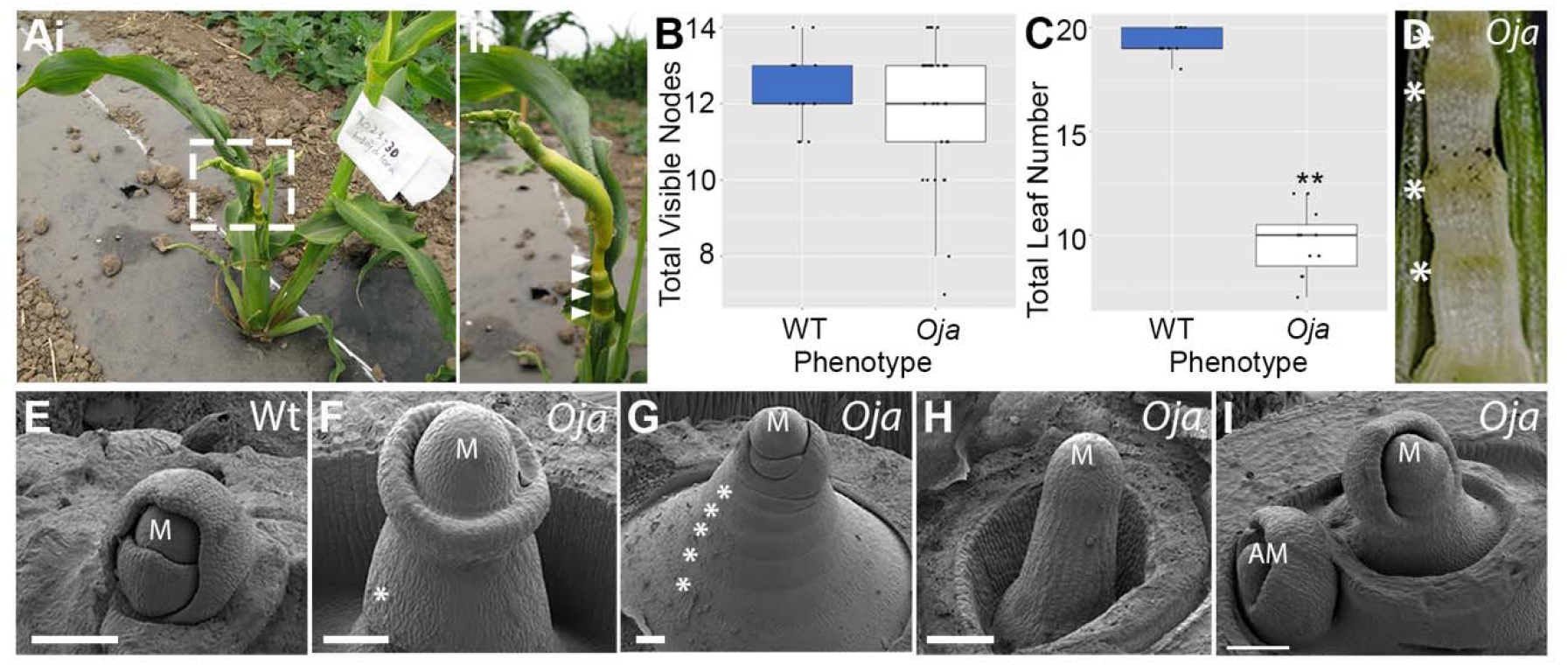
Hoja loca1 mutants initiate new phytomers but fail to initiate leaves. (A-C) Mature *Oja* plants are often short, and despite having a similar number of visible nodes (B, p=0.2537, Welch’s 2-Sample T-Test, df= 39.8, n *Oja*=29, Wt=14), they have fewer leaves (C, ** p= 4.17e-11, Welch’s 2-Sample T-Test, df=14.194, n= 10 of each phenotype) than wild type. In particular, the nodes below the tassel are often bare (Aii, arrowheads). Bare nodes can also be identified in cross-sections through developing shoots (D). The lack of leaf initiation is clear at the earliest stages of development. Scanning electron micrographs (SEMs) of wild type (E) and mutant (F-I) vegetative meristems with attached leaf primordia show severe defects in *Oja*. Some leaf primordia fail to arch over the meristem, instead forming a ring-shaped primordium (F). Some have clear ridges where internodes have formed below the meristem (G), which would normally be concealed by leaves in wild-type plants. Other mutants form pin-shaped meristems (H). Often the tiller buds in *Oja* are normal (I) allowing the plant to produce progeny. The internodes are indicated by the *, meristems by ‘M’, axillary meristems by ‘AM’. Scale bars are 100μm.

To identify whether leaves failed to initiate, or whether they initiated but failed to elaborate, we used scanning electron microscopy (SEM) to examine the earliest primordial stages of leaf development. Wild-type maize leaves at the P_1_ stage are a ring of cells that surround the meristem. At P_2_, the leaf begins to grow in the proximal distal dimension, with growth highest at the midvein position. By P_3_, the leaf primordium covers the top of the meristem and the margins begin to expand (Figure 3E). Further growth of the leaf margins at the P_4_ stage envelopes the younger leaves and meristem (Hay and Hake, 2004; Scanlon et al., 1996). Longitudinal growth of the internode and sheath occur later in development. Compared to wild-type seedlings where developing leaf primordia always obscured the internodes, the internodes were often visible in severe *Oja* shoots (Figure 3F-I). The meristem in Figure 3F has a single ring primordium which may be the early stages of a tube leaf, and below this ring, no older leaves have initiated, leaving an elongated, exposed internode. The meristem in Figure 3G initiated five internodes without leaves, before more leaf primordia were initiated. In some *Oja* mutants, the leaf initiation defect was so severe that the shoot apex ended in pin-like barren meristems as in Figure 3H. In many mutants, axillary meristems formed which contained more normal leaf initiation (Figure 3I). In fact, the mature plant in Figure 3A had a fertile tiller although the main shoot ended in a tube leaf with skipped leaf initiation. The SEM data suggest that the discrepancy between node number and leaf number in *Oja* is due to patterning during the earliest stages of leaf primordium initiation. This finding supports the hypothesis that *Oja* mutants fail to initiate leaves, despite initiating new phytomers.

The skipped leaf initiation also explains some of the seedling architecture differences. For example, the normal seedling in Figure 1A has four visible leaves, while the seedling in Figure 1B has only three leaves visible, with the second leaf the size of a normal third leaf. In addition, the distance between the first and second leaf is much greater than in the wild type. This observation suggests that leaf two was skipped in the mutant.

### *Hoja loca1* encodes the AUX/IAA protein, ZmIAA28

To identify the causal mutation, we mapped the Mo17 *Oja* region in a backcross population to A619. Bulk segregant mapping placed *Oja* on chromosome 8 (Figure 4). SSR primers were used to narrow the position to an interval of 1 Mb containing 19 annotated genes (Figure 4Aiii). Using published RNAseq data we narrowed the candidate gene list further, as only 10 genes in the interval were reported as expressed in the shoot apical meristem or leaf primordia (Figure 4Aiii, green).

**Figure 4.**
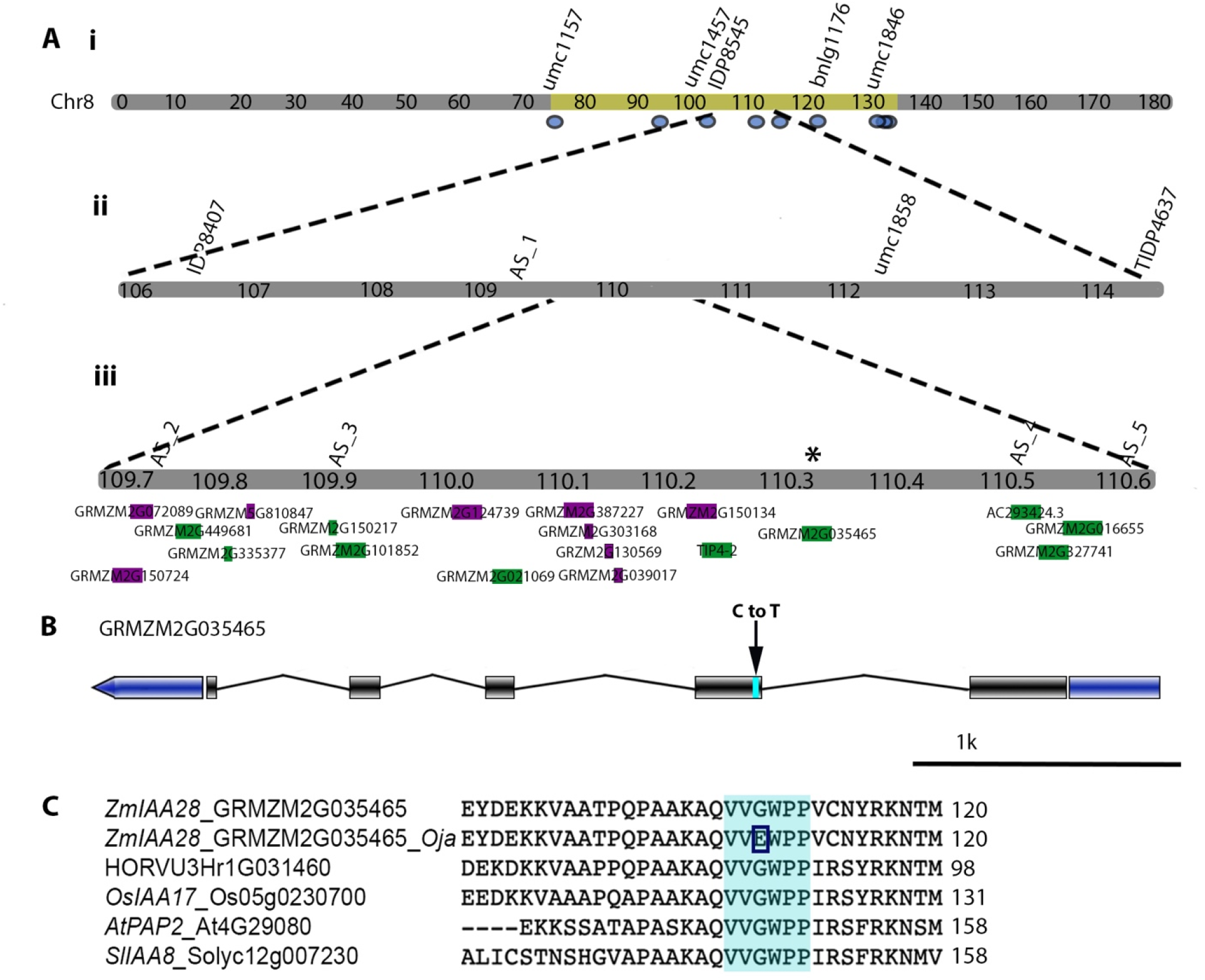
*Hoja loca1* maps to ZmIAA28 on Chr8. Using short sequence repeat (SSR) PCR markers, *Hoja* was mapped to Chr8 between 109.7-110.6Mb (A). Custom SSR markers are labeled as ‘AS#’. Of the 19 genes in the fine mapping interval (Aiii), 10 are expressed in shoot meristems and leaf primordia (green), 9 are not (magenta) based on published RNAseq datasets. Whole genome sequencing (WGS) data identified 9 high to moderate effect SNPs in gene bodies in the larger mapping region (Ai, blue ovals, Chr8:75-135Mb). Of these, only one is an EMS-type SNP that falls within the fine mapping interval (Aiii, *). (B)The SNP causes a C to T change in the second exon of GRMZM2G035465, *ZmIAA28* which results in a glycine (G) to glutamic acid (E) amino acid change in the highly conserved core degron motif (C, cyan box).

To further map the interval, we used whole genome sequencing of two bulk pools, one wild type and one mutant pool, each with 40 individuals. A large interval was detected on chromosome 8 between 75 and 135 Mb. To generate a candidate SNP list, we removed SNPs reported in the HapMapv3 from Panzea (www.panzea.org/data) and limited the reported SNPs to EMS-type G to A, or C to T. Within the large interval, there were nine moderate or high-effect SNPs within annotated gene models (Figure 4Ai), but only one SNP detected within the fine SSR mapping interval (Figure 4Aiii, *). This change was a C to T nucleotide transition in GRMZM2G035465, which encodes ZmIAA28 (Figure 4B). The C to T transition causes a missense mutation from a Glycine to Glutamic Acid in the center of the highly conserved degron motif (Figure 4C). Similar mutations in other AUX/IAA proteins prevent degradation in response to high auxin (Reed, 2001) suggesting that *Oja* is caused by a stabilizing mutation in the degron motif, preventing degradation of ZmIAA28 in the presence of auxin.

### *Hoja loca1* shows dysregulation of auxin signaling

To assess the effect of the *Oja1* mutation on gene expression in vegetative meristem and developing leaf primordia, we sequenced mRNA extracted from dissected meristems with P1/P2 primordia attached from 3-week-old seedlings (Figure 5A). Given the variability in phenotype, each individual was used to make a single library, with 36 libraries sequenced in total. By scoring the libraries based on the seedling leaf phenotype, we classified the libraries into two groups: wild type and mutant (orange and blue annotations indicated on the principal component analysis plot in Figure 5B). Differential gene expression analysis identified 2877 significantly differentially expressed genes (padj<0.05) with a log2FC of >0.59 or <0.59, of which 689 were down in the mutant-scored samples compared to the wild type. GO term analysis showed enrichment for many terms, including those associated with auxin signaling, meristem regulation and leaf development (Figure 5C). Consistent with the phenotype of *Oja*, genes associated with leaf development are down, and those associated with meristem regulation are up in the mutant-scored samples compared to the wild type-scored samples. This suggests that ZmIAA28 has a role in regulation of auxin response, and leaf development.

**Figure 5.**
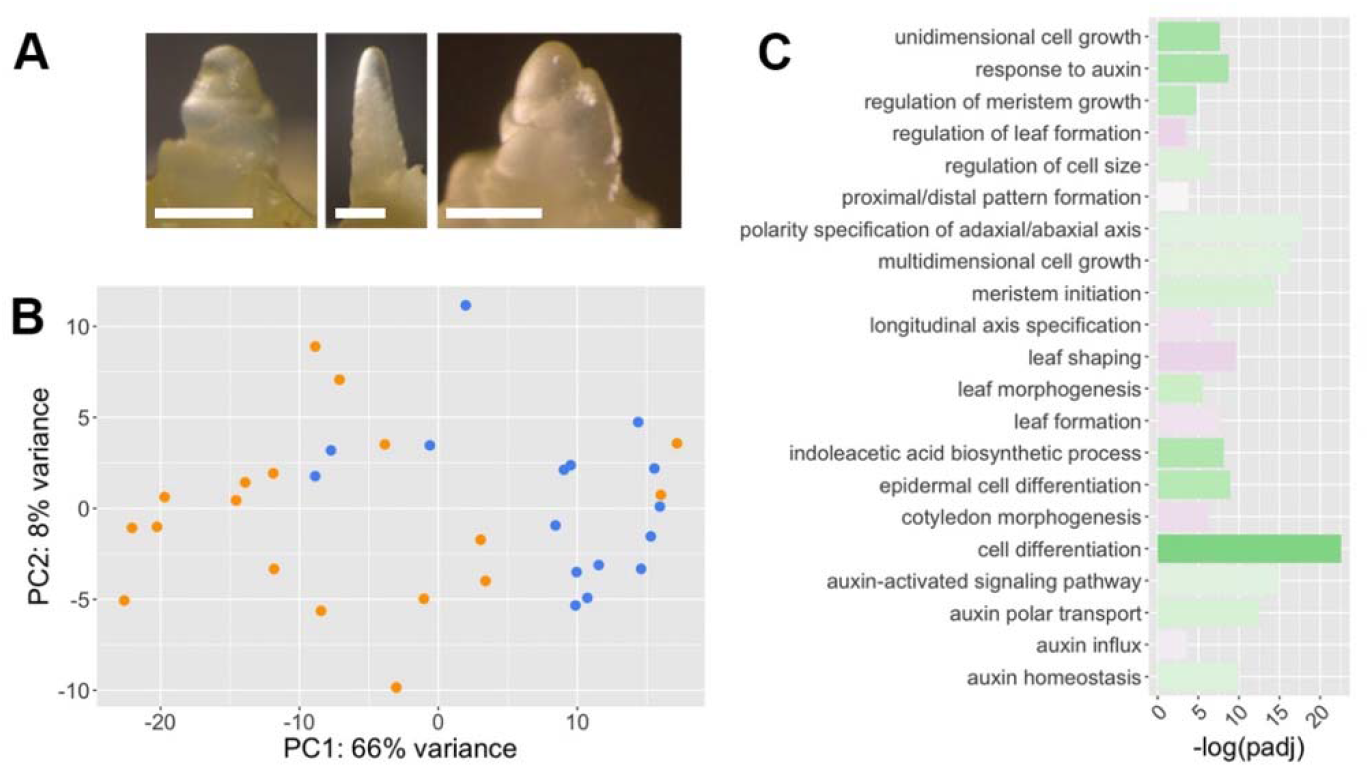
Hoja loca1 mutants show changes in auxin, leaf development, and meristem regulation genes. (A) Examples of the dissected shoot apical meristems harvested for the RNAseq experiment. Scalebar:100μm. (B) Principal component analysis plot (PCA) of all RNAseq libraries, coloured by seeding phenotype at time of harvest; mutant (orange) or wild-type (blue) (n=100). (C) Significantly enriched GO terms (–log(padj) >1) associated with auxin, meristem regulation and leaf development for all genes that are significantly differentially expressed between mutant and wild type scored samples (padj<0.05, log2foldchange >0.59 or <-0.59). Z-score indicates whether the genes in the GO term are on average upregulated (green) or downregulated (purple).

### Transcriptomes of individual *Hoja loca1* meristems and leaf primordia can differentiate four distinct transcriptional profiles, correlated with the four phytomer phenotypes

To further investigate how the different leaf phenotypes of *Oja* may arise, we interrogated the transcriptional profiles of each individual sample. We asked whether we could identify four transcriptional profiles representing the four potential phenotypes of an initiating *Oja* phytomer: normal leaf, midribless leaf, tube leaf and no leaf initiation.

We first filtered the expressed gene list to those shown to be expressed in the meristem and developing leaf primordia based on (Knaeur 2019) to reduce noise caused by differing amounts of stem and internode tissue harvested with each sample. Using the top 1500 most variable of these genes, we carried out hierarchical clustering and identified four clusters using K-means (Figure 6A). Overlaying these clusters on a PCA plot of the top 100 most variable of all expressed genes showed clear separation of the four groups along PC1 (Figure 6B). This indicated that the four clusters (C1-4) were transcriptionally distinct.

**Figure 6.**
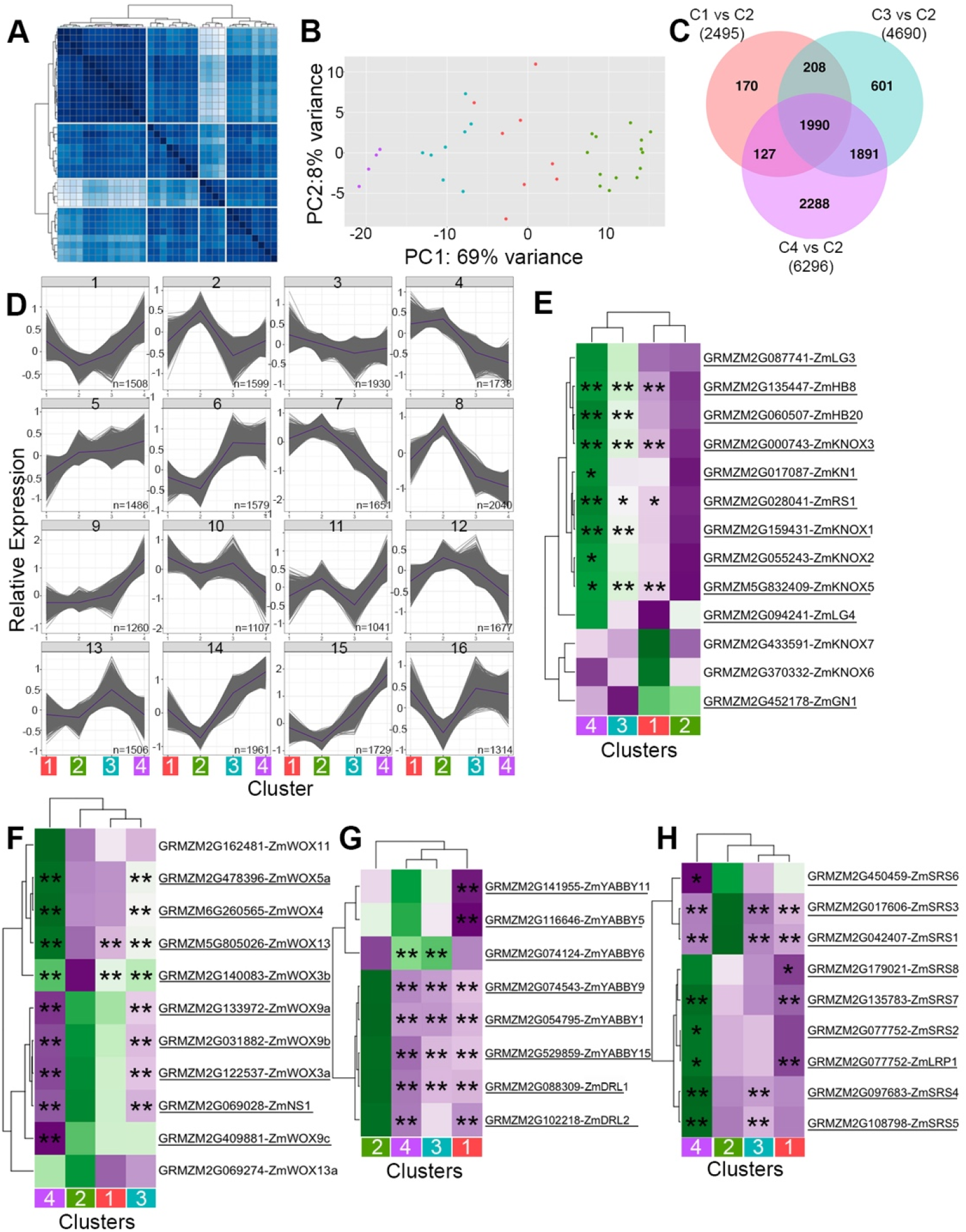
Transcriptional profiles of individual vegetative meristems enable identification of four transcriptional profiles associated with mutant leaf phenotypes. (A) Hierarchical clustering of 1 minus the Pearsons correlation of sample distances (dark blue most similar) based on the top 1500 most variable leaf and meristem associated expressed genes. K=4. (B) Principal component analysis plot (PCA) coloured by the 4 clusters identified in the hierarchical clustering (n=100), cluster 1 (red), cluster 2 (green), cluster 3 (blue), cluster 4 (purple). (C) Venn diagram showing the overlap in significantly differentially expressed (sigDE) genes in comparisons with cluster 2 (C2) which have a log2foldchange of >0.59 or <-0.59. C#vsC# indicates cluster comparison. (#) indicate the total number of sigDE genes, numbers indicate the number of genes in each category. (D) 4×4 self-organizing maps built from all expressed genes (defined as more than 5 rpm in more than 3 samples) in each cluster. Nodes 1-16 are shown. N indicates number of genes in the node. (E-H) Heatmaps of average FPKM values of expressed *KNOX* (E), WOX (F), *YABBY* (G) *and SRS* (H) genes in clusters 1-4. Colouring is relative average FPKM per row, green is positive, purple is negative. Genes that are significantly differentially expressed in one or more pairwise comparisons with cluster 2 are underlined (padj<0.05), “*” indicates genes with a log2foldchange >0.59 or <-0.59, “**” indicate genes with log2foldchange >1.0 or <-1.0.

To assess which cluster may be associated with each phytomer phenotype, we generated self-organizing maps (SOMs) of all expressed genes across C1-4 (Figure 6D) and annotated the nodes with genes involved in leaf initiation and development. A defining feature of initiating leaf primordia is the specific downregulation of *KNOX* genes in the P_0_ and the activation of leaf patterning genes. For the four phytomer phenotypes; normal leaf, midribless leaf, tube leaf and no leaf initiation; we predicted that these genes would show distinct profiles. In phytomers initiating normal leaves we would expect low *KNOX* gene expression, with high leaf patterning gene expression, and the converse for phytomers failing to initiate leaves. Phytomers initiating midribless leaves and tubes leaves would have an intermediate profile. We found that most *KNOX* genes were in node 14, (Figure 6D) where expression was at the lowest level in C2 (Figure 6E), consistent with *KNOX* genes being downregulated during normal leaf initiation. Genes associated with leaf development were found in nodes where expression was highest in C2 (e.g. node 7 contains *CUP-SHAPED-COTYLEDON* genes and *WUSCHEL-LIKE-HOMEOBOX* genes known to be involved in leaf development and organ initiation). This supports the hypothesis that C2 represents meristems initiating phytomers with normal leaves.

Conversely, higher *KNOX* (node 14) and lower leaf development gene expression (e.g. nodes 7 and 8) associated with C4 (Figure 6D). This is consistent with the hypothesis that C4 represents phytomers with a ‘no leaf initiation’ phenotype. This is further supported by the expectation that phytomers failing to initiate leaves will be most transcriptionally distinct from those initiating normal leaves, and the PCA and hierarchical clustering both show C4 as most different to C2 (Figure 6B). Combined, these data suggests that C2 represents phytomers initiating normal leaves, and C4 represents phytomers with no leaf initiation.

To identify which of the remaining two clusters may correlate with either the tube leaf or midribless leaf phytomer phenotype we analysed the expression profiles of medio-lateral leaf patterning genes. We predicted that these genes would be misregulated in phytomers initiating midribless or tube leaves, and be downregulated in C4 as these phytomers are not initiating leaves.

Tube leaves lack leaf margins, and this should be reflected by reduced expression of genes like the *WOX* gene *NARROWSHEATH1 (NS1)* which defines the leaf margin (Nardmann et al., 2004; Scanlon and Freeling, 1997). Many *WOX* genes, including *NS1*, are found in node 7 where expression is lowest in C3 and C4. Low *WOX* expression in C3 could indicate that this group represents phytomers initiating tube-shaped leaves.

Few genes have been identified as responsible for midribless identity, but the *YABBY* genes *DROPPING LEAF 1* (*DRL1/IFA1*) and 2 (*DRL2*)(Strable et al., 2017) both have defects in midrib development. We would predict that genes involved in midrib development would be downregulated in phytomers initiating midribless leaves. *DRL1* and *DRL2* are located in node 8, where expression is highest in C2 and low in all other clusters, making it difficult to differentiate between C1 and C3. Given that the midribless leaf phenotype appears most similar to a normal leaf, we predict that the cluster most similar to C2 would represent phytomers initiating midribless leaves. Based on the PCA and the differential gene expression analysis, C1 was most similar in transcriptional profile to C2 (Figure 6B,C, pink). We therefore hypothesise that C1 represents phytomers initiating midribless leaves, and C3 represents phytomers initiating tube leaves.

### Each phytomer phenotype can be associated with specific gene expression changes

To further test the hypothesis that each cluster represents a different phytomer phenotype (C1-midribless leaf, C2-normal leaf, C3-tube leaf, C4-no leaf initiation), we carried out differential gene expression analysis for each ‘mutant’ cluster (C1, 3 and 4) in comparison with the hypothesised normal leaf representative (C2). Overlap of the significantly differentially expressed gene lists (padj <0.05, log2FC<-0.59 or>0.59, Figure 6C, E-H) showed distinct transcriptional profiles for each cluster, suggesting that each phytomer phenotype may be underpinned by different gene expression changes triggered by variable effects of ZmIAA28 stabilization on auxin signaling.

To identify genes potentially important for each phenotype we overlaid the significantly differentially expressed genes (log2 fold change >0.59 or <-0.59) for each comparison, and identified 170 genes specific to C1, 601 to C3, and 2288 to C4. C4 showed the most unique significantly differentially expressed genes, many of which are important components of leaf initiation. These genes included the boundary genes *CUC2* and *CUC3*, which were significantly downregulated, consistent with their roles in specification of the boundary between the primordium and meristem (Hibara et al., 2006; Aida et al., 1999). This further supports the hypothesis that C4 represents phytomers which fail to initiate leaves, and suggests ZmIAA28 degradation in the presence of auxin is involved in triggering leaf initiation.

We predict that C1 and C3 represent phytomers initiating leaves with medial-lateral patterning defects. *WOX* genes exhibited distinct gene expression profiles across the clusters, with opposite expression profiles in C3 compared to C2. *WOX3b, 4, 5a* and *13* were significantly upregulated in C3, whereas *WOX3a, 9a, 9b,* and *NS1* were all significantly downregulated (Figure 6F). This finding suggests that medial-lateral patterning and marginal domain development is severely perturbed in C3, correlating with the tube leaf phenotype. The analysis demonstrates that auxin signalling via ZmIAA28 degradation is important in regulation of *WOX* gene expression for medial-lateral domain patterning.

Midribless leaves show defects in midrib formation and in vein distribution in the leaf blade. We therefore analysed the expression profiles of genes with potential roles in midrib and vein patterning. *YABBY1, 9*, *15, DRL1 and DRL2* are all significantly downregulated in C1 compared to C2 (Figure 6G). This finding suggests that ZmIAA28 may link auxin signalling with midrib formation via regulation of DRL1 and 2. C1 also showed significant downregulation of multiple *SHORT INTERNODES (SHI) / STYLISH (STY) / SHI RELATED SEQUENCE (SRS)* genes (Figure 6H), which in Arabidopsis are involved in vascular patterning (Baylis et al., 2013) and have been shown to be regulated by AUX/IAA proteins (Singh et al., 2020). ZmIAA28 may therefore also be involved in auxin regulation of *SRS* gene expression during grass leaf vascular patterning.

### *ZmIAA28* is involved in the auxin maxima maintenance preceding leaf initiation

The no leaf initiation phenotype of *Oja* led us to ask if we could use *Oja* to dissect the link between organ initiation and auxin maxima formation. Using the comparison between C4 and C2, we first interrogated the expression profile of genes known to be involved in organ initiation. We then used *in situ* hybridisation and immunolocalisation to test gene expression in a spatial manner.

In C4, *KNOX* genes are significantly upregulated, indicating that *KNOX* gene downregulation is aberrant in *Oja* meristems in which leaf initiation fails. We used antibodies to KNOTTED1 (KN1) (Smith et al., 1992) in order to follow leaf initiation defects in the meristem. KN1 accumulates in the meristem and unexpanded stem and is clearly down-regulated at the P_0_ region, remaining absent in normal maize leaves (Figure 7A). In *Oja* mutants we see strong accumulation at the tip of the meristem but, unlike the normal meristem, KN1 down-regulation appears on both sides of the meristem, suggesting that this P_0_ encircles the meristem (Figure 7B). The encircling downregulation could be indicative of tube leaf initiation. This particular meristem has also failed to initiate at least two leaves, and KN1 localisation is still maintained in the internode region below the meristem (Figure 7B). This variable downregulation of *KNOX* genes supports the hypothesis that ZmIAA28 may be upstream of the downregulation of *KNOX* genes in initiating primordia.

**Figure 7.**
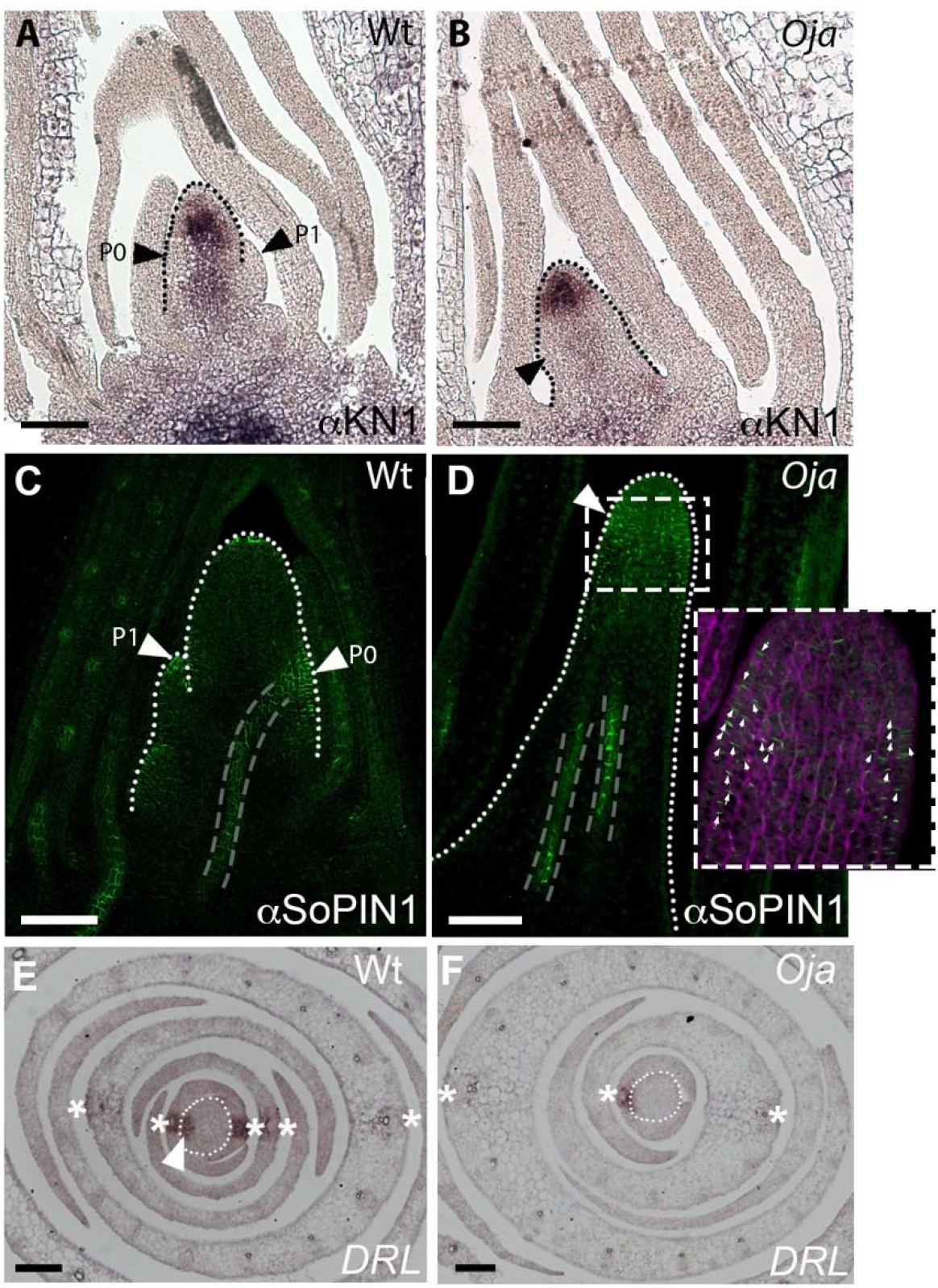
Hoja loca1 have defects in organ initiation markers localization. KNOTTED1 (KN1) immunolocalisation (dark brown/black, A-B) and SISTER OF PINFORMED1 (SoPIN1, green, C-D) in longitudinal sections through 3-week-old seedlings in WT (A,C) and mutant (B,D) plants. The inset in D shows a magnified image of the boxed region, the arrowheads indicate the orientation of the SoPIN1 localization (green) within the cell outline (magenta). Large arrowheads indicate the P_0_ and P_1_ domains. Dotted lines indicate the outline of the meristem region. The white dotted line in A indicate the P_0_. Grey dashed lines indicate vascular traces. (E-F) *In situ* hybridization of *drooping leaf* (dark brown, DL) in transverse cross-sections through 3-week-old seedlings, in wild type (E) and mutant (F) plants. Sections are 100μm from the tip of the meristem. * indicate the position of the midvein in successive leaves, the white dotted line outlines the meristem. Scale bars are 100μm.

Preceding *KNOX* downregulation, the auxin efflux transporter PIN1 moves auxin in the epidermis to the site of leaf initiation, where an auxin maxima forms. From the P_0_, PIN1 proteins move auxin toward pre-existing vasculature, predicting the path of provascular traces (Reinhardt et al., 2003). In many species outside of the *Brassica*, PIN1 proteins have duplicated and diversified function. Sister of PIN1 (SoPIN1) localizes at the leaf initiation site and is also detected in vasculature in grasses (O’Connor et al., 2014). PIN1a and PIN1b have roles in vascular formation and stem growth (O’Connor et al., 2017). We found that all PIN gene expression was significantly downregulated in C4 compared to C2, suggesting that the feedback mechanism that maintains auxin maxima during organ initiation may be defective in *Oja*. As SoPIN1 is essential for organ initiation in the grasses (O’Connor et al., 2017) and is redundantly involved in vascular development, we chose to focus on SoPIN1 accumulation in the plant. To determine whether SoPIN1 patterning was defective in *Oja* meristems, we used an antibody to SoPIN1 to detect protein accumulation at the P_0_ site and in the developing pre-vascular strand in wild-type plants (Figure 7C). In wild-type meristems, SoPIN1 convergence points form in the epidermis of the meristem at the P_0_, which persists through to the tip of the P_1_ leaf (Figure 7C). Strong internal traces of SoPIN1 are also observed highlighting the developing vascular strands that link the primordium apex to the stem vascular system (Figure 7C). In the *Oja* mutant shown, which has a number of skipped leaf initiations and appears as a pin, the SoPIN1 localization is throughout the tip of the meristem rather than localizing strongly to the P_0_ region (Figure 7D). Normal accumulation and polarization can be seen in a vein trace within the mutant meristem, but this does not connect to the epidermis. This result suggests that SoPIN1 can polarize and be coordinated across the tissue. However, SoPIN1 fails to form a strong convergence point needed at the P_0_ in *Oja*, implying that ZmIAA28 may be involved in the feedback regulation of SoPIN1 at lateral organ initiation sites, essential for organ initiation. In support of this hypothesis we observe changes in the expression of several genes involved in PIN protein localisation, including the PINOID protein BIF2 (McSteen and Hake, 2001), which is significantly downregulated in C4 compared to C2.

Downstream of auxin maxima formation and *KNOX* gene downregulation is activation of leaf specific genes, such as *DRL1* and *2* (Strable et al., 2017). DRL genes are early markers of leaf patterning and are expressed at the P_0_ and at the midrib in leaf primordia. To explore the effect of *Oja* on *DRL* expression, we used *in situ* hybridization. In the normal seedling, expression was detected in the P_0_ and each subsequent midvein region (Figure 7E). In the *Oja* mutant shown, *DRL* expression was similar at the midvein region in normal developing leaves, however, there was no detectable expression in the P_0_ and fewer leaves had initiated. As DRL is involved in midrib specification (Strable et al., 2017), this lack of DRL in the P_0_ could be due to the lack of leaf initiation or due to the formation of a midribless leaf. Nevertheless, as an early marker of leaf patterning this expression pattern supports the hypothesis that *Oja* mutants fail to specify the P_0_ leaf correctly despite defining the rest of the phytomer.

In summary, these findings suggest that *Oja* has a defect in leaf initiation and patterning, not just leaf outgrowth, which leads to the reduced leaf number compared to node number.

## DISCUSSION

The dominant *Hoja loca1* (*Oja*) mutant, caused by a mutation in ZmIAA28, specifically affects aerial organ initiation and medial/lateral patterning in the leaf, illustrating highly specific AUX/IAA function essential for the fine-tuning of plant development by auxin. By analysing individual meristems, we were able to use the variable phytomer phenotypes of *Oja* to dissect potential regulatory roles of ZmIAA28 that may be responsive to different auxin concentrations.

AUX/IAA proteins function as repressors in low auxin conditions by interacting with ARF transcription factors and preventing their activation of auxin responsive genes. An increase in auxin promotes an interaction between AUX/IAA and Transport Inhibitor Response1 (TIR1), an F-box component of the SCF complex that targets proteins for degradation. Binding of AUX/IAA to TIR1 in the presence of auxin leads to AUX/IAA degradation and thus releases ARF repression (Dharmasiri et al., 2005; Kepinski and Leyser, 2005). As the AUX/IAA gene family is large, redundancy has mostly hidden loss-of-function phenotypes (Overvoorde et al., 2005). Instead, genetic analysis has identified dominant AUX/IAA mutations that stabilize these normally short-lived proteins by disrupting their proteolysis or reducing their ability to bind to TIR1 (Ouellet et al., 2001; Ramos et al., 2001; Tian et al., 2003; Worley et al., 2000; Yang et al., 2004). Most mutations, including *Oja*, are located in domain II in the highly conserved core degron motif (Fukaki et al., 2002; Nagpal et al., 2000; Rogg et al., 2001; Rouse et al., 1998; Zenser et al., 2001) which interacts with TIR1 in the presence of auxin (Tan et al., 2007).

Nine clades of AUX/IAA proteins have been identified (Matthes et al., 2019), allowing for significant subfunctionalization during the evolution of the AUX/IAA family. The majority of *Arabidopsis* AUX/IAA mutants were first described by their agravitropic root, reduced lateral roots, and/or shorter root phenotypes (Fukaki et al., 2002; Nagpal et al., 2000; Rouse et al., 1998). Mutants that were identified by a shoot phenotype such as *bodenlos*, *Shy2*-2 and *axr3* have pleiotropic phenotypes, also affecting the roots. For example, the semi-dominant *Shy2* mutant was identified as a suppressor of a phytochrome elongated hypocotyl phenotype but also has root defects and curled leaves (Reed et al., 1998; Tian and Reed, 1999). Embryonic patterning defects were identified in *bodenlos* homozygotes but mature heterozygous plants also have root defects (Hamann et al., 2002). *Axr3* mutants develop curled leaves (Leyser et al., 1996; Pérez-Pérez et al., 2010) due to defects in coordination between the abaxial and adaxial surfaces of the leaf. When these mutant phenotypes are mapped to the AUX/IAA clades, no obvious pattern is observed, suggesting that function is not conserved within a clade. This finding highlights the importance of studying the specific function of AUX/IAA proteins in multiple species, as genetic similarity is unlikely to correspond to related functionality.

Our work suggests that ZmIAA28 acts specifically during aerial organ initiation, potentially upstream of *KNOX* downregulation. This contrasts with the function of the tomato class VIII AUX/IAA, SlIAA9. Tomato leaf margin complexity is dependent on the ectopic expression of *KNOX* genes in the leaf lamina. However, reduction of SlIAA9 expression causes a loss of leaf margin complexity, independent of *KNOX* gene expression as determined by RT-PCR (Wang et al., 2005). This suggests that although ZmIAA28 and SlIAA9 are in the same clade, they differ in their relationship with *KNOX* gene regulation. This further argues for the importance for studying specific AUX/IAA functions in different species.

AUX/IAA mutants in maize appear to be less pleiotropic than *Arabidopsis*; their defects are confined to specific parts of the plant. For example, the *rootless with undetectable meristem* (*rum1*) mutant is deficient in primary and lateral root initiation with no defects in the aerial parts (Woll et al., 2005). *rum1* encodes *ZmIAA10* and, unlike other described AUX/IAA mutants that have a missense mutation in the degron, *rum1* carries a deletion that includes the GWPPV degron sequence in domain II (Von Behrens et al., 2011). The *Bif1* (ZmIAA27) and *Bif4* (ZmIAA20) AUX/IAA mutants reduce the branching in male and female inflorescences (Barazesh et al., 2009; Galli et al., 2015), however there is no effect on vegetative growth (Galli et al., 2015). *Oja* mutants have striking defects in leaf development but have normal roots and flowers if they make inflorescences. This suggests that ZmIAA10, ZmIAA20, ZmIAA27 and ZmIAA28 have tissue-specific functions. These specificities do not appear to arise from tissue specific expression as, based on published RNAseq data, ZmIAA27 and ZmIAA28 have similar expression patterns. *RUM1* is expressed at very low levels compared to a close paralog which is expressed at high levels in many root tissues as well as the leaf (Zhang et al., 2016). Perhaps the tissue specificity results from the interaction of each AUX/IAA with different ARFs or different TIR proteins and these are responsible for the observed phenotypes in the dominant mutants (Salehin et al., 2015).

Through investigating leaf initiation, we found that SoPIN1 failed to converge at the P_0_ in *Oja* mutants suggesting that ZmIAA28 plays a role in the feedback signaling required for forming the convergence point that triggers organ initiation. A role for the TIR/ARF/AUX/IAA module in regulating cellular polarisation has also been observed in *Arabidopsis* in which induction of the ARF, *MONOPTEROS* (MP) in the *mp* mutant background triggered spontaneous PIN1 convergence point formation and primordium outgrowth in the normally barren *mp* meristem (Bhatia et al., 2016). Interestingly, although leaf initiation was severely affected in *Oja*, many other organ initiation events occurred normally. For example, ligule development requires reactivation of the organ initiation pathway (Johnston et al., 2014) but *Oja* mutants have no ligule defect, suggesting that different AUX/IAAs may be involved in ligule development.

Given that formation of the SoPIN1 convergence point at the P_0_ stimulates midvein development, it is likely that ZmIAA28 is required for the signaling that promotes midrib formation at the midvein, influencing medio-lateral patterning and possibly subsequent *DRL* expression. Supporting the role of auxin signaling upstream of *DRL* expression, DAPseq analyses in maize have identified ARF binding sites upstream of both maize *DRL* genes (Galli et al., 2018). In midribless *Oja* mutant leaves, the midrib domain appears to be completely lost, which is more severe than in the *drl* double mutant (Strable et al., 2017), suggesting that other factors, also influenced by auxin signaling, are required to specify midrib differentiation in addition to DRL.

Mediolateral development in the leaf is driven by *WOX* gene activity (Conklin et al., 2021; Zhang et al., 2020). Our transcriptional analyses highlight wide-spread miss-regulation of *WOX* gene expression in *Oja*, with most significant downregulation associated with potential tube leaf formation, and failure of leaf initiation. The formation of tube-shaped leaves in maize can be induced by incubation with the auxin inhibitor NPA (Scanlon, 2003) which leads to a failure of sheath margin separation, and this correlates with a high accumulation of ZmPIN1a in this region at the P4 stage of leaf development (Johnston et al., 2015). Recent work has also shown that NS1 is induced by auxin treatment (Conklin et al., 2021). This suggests that ZmIAA28 may be involved in translation of the auxin concentration signal to leaf margin specification and later margin development.

Analysis of proximal-distal patterning mutants gave rise to the hypothesis that the midrib domain acts as a signaling centre to coordinate ligule development from the midrib to the leaf margins (Becraft and Freeling, 1991; Harper and Freeling, 1996; Moon et al., 2013). However, proximal-distal patterning is normal in *Oja* leaves lacking the midrib. This suggests that it is not the midrib domain itself that links medial-lateral and proximal-distal patterning but potentially medial-lateral patterning factors that are largely independent of ZmIAA28 regulation of midrib formation.

The variability of the *Oja* phenotype is not unusual, having also been observed in other AUX/IAA mutants such as in the *Arabidopsis* mutant *bodenlos,* with some mutants missing hypocotyl and root and others just missing a root (Hamann et al., 2002). These variable phenotypes could arise from stochastic over- and under-compensation of the auxin signaling network. The sensitivity of different tissues to these stochastic changes in auxin concentration could arise through different combinations of TIRs, ARFs and other auxin signaling components (Powers and Strader, 2020; Parry et al., 2009; Havens et al., 2012). In such a scenario, the four clusters of gene expression and corresponding phytomer phenotypes identified in this work, may represent stable end-states that arise from perturbations of the auxin pathway very early in development. As such, when ZmIAA28 is not degraded in the presence of high auxin, as it is in wild type, the three mutant phytomer phenotypes result from sub-optimal compensatory mechanisms cascading to islands of stability. *Oja* therefore reveals three regions where response to high auxin concentrations during the earliest phases of leaf development is crucial in defining leaf shape: initiation, formation of a midrib/ vascular patterning, and separation of the leaf margins.

The phytomer consists of a node, internode, axillary bud and organ. To maintain correct plant architecture, the development of each component is tightly regulated. *Oja* is unusual in that this coordination within the phytomer unit is apparently broken, allowing the initiation of nodes and internodes without accompanying organs. This finding suggests that auxin is not only involved in the initiation of organs, but also in the regulation of phytomer development as a whole. The *Oja* mutant illustrates the species-specific subfunctionalisation of the AUX/IAA proteins in regulating plant development. In contrast to other known AUX/IAAs, ZmIAA28 specifically affects initiation of aerial organs and impacts medio-lateral patterning. As further studies identify functions of other grass AUX/IAAs it will be of interest to compare them to *Oja* to see how far this subfunctionalisation holds. This will be particularly interesting to compare to *Arabidopsis aux/iaa* mutants which appear to be more pleiotropic, and to determine the reasons for the species-specific differences.

## Methods

### Isolation of *Hoja loca1*

*Hoja loca1* (*Oja*) was discovered by a forward genetic approach. Pollen of the Mo17 maize inbred was treated with ethyl methanesulfonate (EMS) and crossed onto the B73 inbred. Hybrid F1 seed were planted in the field and screened. The single mutant plant was open pollinated and crossed to multiple inbreds the following season. Further crosses established the dominance of the mutant phenotype although often less than 50% of the plants showed the phenotype, indicating low phenotypic penetrance.

Each generation displayed unpredictable variability in phenotypic expression ranging from mild to severe. Crosses made with mildly affected plants gave rise to severe plants and vice versa. Due to the severity of the phenotype, homozygotes were rarely obtained, thus the observations are exclusively with heterozygotes. Analysis was carried out in plants with at least four backcrosses to B73.

### Plant Growth Conditions

Analyses were carried out in families segregating for *Hoja loca1* after at least four backcrosses to the B73 inbred line. Plants were grown in Supersoil Brand Potting Soil (composted bark (60-85%), Sawdust (60-85%), Sphagnum Peat Moss (10-30%)) in standard greenhouse conditions. Seedlings were grown for 3 weeks in 30 by 45 by 2.5cm trays, and adult plants were grown in 12.5L pots until tassel emergence. Seedlings for the RNAseq analysis were the offspring of a cross between a normal sibling and a mutant *Oja* sibling, grown in M3 soil mix with insecticide in large trays equally spaced, in the Grodome facility at the University of Edinburgh with supplementary lighting day/night cycles of 21C, 16hours / 15C, 8 hours.

### Root Analysis

Seeds were rehydrated overnight in sterile water, then sterilized with 10% bleach for 20 minutes, before washing with sterile water. Seeds were then germinated on damp chromotography paper in vertically fixed plastic wallets. After one week, the seedlings were photographed, then rotated 90°. The seedlings were left to grow for 24 hours and imaged every hour, for 12 hours. After 24 hours, the final root phenotype was imaged. Primary root length was analysed using FIJI (Schindelin et al., 2012) and crown root number was counted. The seedlings were transplanted to soil and three weeks after germination the shoot phenotype was recorded.

### Histology Plastic Sections

10μm plastic sections were made with Technovit® 9100 from Electron Microscopy Services (Cat. #14655) according to manufacturer instructions. Sections were mounted on glass slides and stained with toluidine blue. Imaging was performed on a Zeiss Axiovert inverted compound microscope with transmitted light.

### Scanning Electron Microscopy

SEMs were performed with material fixed in FAA (3.7% formaldehyde, 50% ethanol, 5% acetic acid) overnight at 4°C and dehydrated in ethanol series to 100%. Samples were critical point dried, sputter coated with palladium, and imaged on a Hitachi S-4700 with 2kV accelerating voltage.

### Tissue for Immunolocalisation and *In Situ* Hybridisation

Plant tissue was collected from 3-week-old seedling shoot apices, dissected and fixed overnight in 4% paraformaldehyde/ 0.1% DMSO/ 0.1% Triton-X100. Fixed samples were transferred through an ethanol and histoclear series before embedding in paraplast plus (McCormick Scientific #19216). 10μm sections of fixed tissue were mounted on probe-on-probe-plus slides (FisherBrand #22-230-900) and dried overnight on a 37°C hotplate before use in either immunolocalisation or *in situ* hybridization.

### Immunolocalisation

Primary antibodies were guinea-pig anti-SoPIN1 as used in (Richardson et al., 2016) and rabbit anti-KNOTTED1 (KN1) as used in (Bolduc et al., 2012b). Secondary antibodies were commercial anti-guinea pig conjugated to Alexa-fluor594 (Jackson ImmunoResearch, #706-585-148) and anti-rabbit alkaline phosphatase conjugate (Jackson ImmunoResearch, #111-055-045). All antibodies were used at a 1:200 dilution in 1% bovine serum albumin (BSA) in PBS.

Immunolocalisation used a protocol based on (Conti and Bradley, 2007), and was as follows. Slides were de-paraffinised using histoclear, then rehydrated through an ethanol series to water. Samples were then boiled in 10mM sodium citrate, pH6, for 10 minutes to retrieve the antigens. Blocking was carried out in 1% BSA/PBS/0.3% Triton-X100 for 3hours. Slides were incubated overnight in primary antibody, before washing in PBS/0.3% Triton-X100. They were then incubated with secondary antibody at room-temperature for 2 hours.

Tissues incubated with anti-KN1 were incubated in a 1:50 dilution of NBT/BCIP (Roche, #11681451001) in 0.05M MgCl_2_/TBS, pH9.5 until a dark precipitate was observed. These slides were then imaged on a Leica dissecting microscope in water.

Tissues incubated with anti-SoPIN1 were stained for 20 minutes in 0.1% calcofluor (fluorescent brightener 28, Sigma-Aldrich #F3543), washed and mounted in water and imaged on a Leica SP8 confocal microscope. The subcellular localization of SoPIN1 was assessed in relation to the calcofluor cell wall stain using FIJI (Schindelin et al., 2012).

### *In situ* Hybridisation

Antisense probes targeted to *DROOPLING LEAF1* (GRMZM2G168893) mRNA were designed as used in (Strable et al., 2017). *In situ* hybridization was carried out as in (Coen et al., 1990) outlined here in brief. Tissue was deparaffinised using histoclear, then rehydrated through an ethanol series. Samples were digested using 0.125mg/mL pronase (Sigma-Aldrich #P6611) in 50mMTris/5mM EDTA, pH7.5, for 15minutes at room temperature. Digestion was stopped by the addition of 0.2% glycine before washing and re-fixing in 4% PFA/PBS for 10minutes. Slides were then treated in 0.1M Triethanolamine-HCl / 0.5% Acetic Anhydride (Sigma-Aldrich #320102) for 10 minutes before dehydrating through an ethanol series. Tissue was incubated overnight with the probes at a 1:50 dilution in hybridisation buffer (0.375M NaCl, 12.5mM Tris-HCl pH8, 12.5mM Sodium Phosphate pH6.8, 6.25mM EDTA, 50% deionized formamide, 12.5% dextran sulfate, 1.25x Denhardt solution, 0.0125mg/mL tRNA) at 50°C. Slides were washed in 0.2% SSC at 50°C, then treated with RNAseA (in 0.5M NaCl/10mM Tris/ 1mM EDTA, pH7.5) at 37°C for 15 mins, before repeating the SSC washes. Slides were blocked in Roche Blocking Reagent (#11-096-176-001) for 30 minutes, washed in1% BSA/0.3 % Triton-X100/TBS, and incubated at room-temperature with a 1:500 dilution of anti-Dioxigenin-AP Fab fragments (Roche, #11093274910) in 1% BSA/0.3 % Triton-X100/TBS for 1hour. Slides were then washed with 1% BSA/0.3 % Triton-X100/TBS, then in 0.05M MgCl_2_/TBS, pH9.5. To visualize the probe localisation, the tissue was incubated in 1:50 dilution of NBT/BCIP in 0.05M MgCl_2_/TBS, pH9.5, until dark precipitate formed. The staining reaction was stopped by transferring to water and the slides were imaged on a Leica MZ16F microscope with a Qimaging Micropublisher camera under water and brightfield conditions.

### Mapping with SSRs

The *Oja* mutation was mapped to chromosome 3 using short sequence repeat (SSR) PCR mapping. Known SSRs were identified using Maize gdb and are listed in Figure 3. Custom SSRs were identified by using the Gramene SSR prediction tool (Temnykh et al., 2001), and specific primers were designed to amplify them.

### Whole Genome Sequencing

10 mm^2^ area leaf discs were collected from 30 wild type and 30 mutant siblings in an advanced B73 introgressed population. Genomic DNA was extracted using a UREA based extraction method (Chen and Dellaporta, 1994) which is as follows. Tissue was ground in UREA extraction buffer (4M urea, 4M NaCl, 1M Tris pH8, 0.5M EDTA, 34mM n-lauroyl sarcosine). Then an equal volume of phenol:chloroform:isoamyl alcohol (25:24:1), was added and vortexed to mix. After centrifugation, the supernatant was decanted and mixed with an equal volume of chloroform. After centrifugation, the supernatant was decanted and mixed with 1/10^th^ volume of 4.4 M NH_4_OAc pH 5.2 and 0.7 volume of isopropanol. Strands of DNA were then collected and washed with 70% ethanol in a separate tube. The dried pellet was resuspended in 100ul of TE.

DNA was quantified using Qubit dsDNA HS Assay Kit (Invitrogen, Q32854), and sent to Novogene for library preparation and sequencing (350bp insert, 150PE). Sequencing was at 10 times coverage of the maize genome. Sequence analysis was carried out using the CYVERSE Atmosphere computing environment. The sequence data was quality checked and the adapters trimmed using fastp, then aligned to the B73_RefGen_v3 maize genome using bowtie2, variants were called using samtools mileup. The variant list was filtered using R, by removing variants recorded in the Panzea HapMapv3 (https://hapmap.ncbi.nlm.nih.gov/) and those present in the wild type pool as well as restricting to EMS type SNPs. Moderate and high effect SNPs specific to the mutant were calculated using SNPEff2 with the B73 RefGenv3 5b+ gene models.

### RNASeq

Seedlings were imaged using a Canon EOS250D camera, then dissected and imaged using a Leica MZ16F stereoscope with a Canon EOS250D camera attached via an LM-Scope DSLR adapter. Seedlings were dissected at 21 and 28 days down to the meristem and the P1/ P2 primordium, and individually flash frozen in liquid nitrogen. RNA was extracted from each individual using TRIzol (Invitrogen), precipitated (0.5M NaCl + 2-propyl alcohol), and washed with 70% ethanol, then resuspended in nuclease-free water. RNA was quantified using a QuBit BR-RNA kit. RNA of sufficient quality was sent to Novogene for 150bp paired-end sequencing, and 36 libraries passed quality checks.

The CYVERSE Atmosphere virtual computing environment was used to carry out sequence quality checks, genome alignments and count extraction. (4CPUs, 32G RAM, with a ubuntu operating system). Sequenced libraries were checked for quality and adapters were trimmed using fastp (Chen et al., 2018), then aligned to the B73 RefGenv3 genome using HiSAT2 (Kim et al., 2019). Counts were called using FeatureCounts (Liao et al., 2014). All downstream analysis was carried out locally (3.6GHz 8-Core Intel Core i9 processor, 32GB RAM) using R packages combined with Jupyter notebooks. Differential gene expression analysis was carried out using DESeq2 (Love et al., 2014) using all genes defined as expressed (in more than 3 samples, with more than 5 counts). Significantly expressed genes were defined as those with a padj value of <0.05, and filtered to a log2 foldchange value of <-0.59 or >0.59. GO term analysis used GOSeq (Young et al., 2010) with the maize-GAMER v3 annotations (Wimalanathan et al., 2018), and significantly enriched GO terms had a Benjamini and Hochberg cutoff padj value of <0.05.

To carry out hierarchical clustering we stringently filtered the expressed gene list to those genes with >5 counts in >10 samples, then filtered for those genes listed as expressed in the meristem and leaf primordia from (Knauer et al., 2019). The top 1500 most variable of these genes were normalised and used to plot the 1 minus the absolute value of the Pearson correlation matrix, and the resulting plot was clustered hierarchically and cut using Kmeans =4. The samples in each cluster were then labelled with cluster identity.

To generate self-organising maps, expressed genes (in more than 3 samples, with more than 5 counts) were normalised, and the average variance stablised value was calculated for each cluster. 4 x 4 node SOM grids were then generated using the kohonen package (Wehrens and Kruisselbrink, 2018).

### Vein Analysis

Cross-sections through the third leaf of 3-week-old seedlings, 1.5cm above the ligule were taken and imaged under water on a Leica DM4000 microscope. Cross-section images were analysed using FIJI (Schindelin et al., 2012). The width of the leaf was recorded, then the number of lateral and intermediate veins were counted. Vein density was calculated as the total number of veins (lateral + intermediate veins) over the width of the leaf.

Vein distribution was assessed by counting the number of intermediate veins between lateral veins. To align the wild type and midribless leaves, the lateral vein closest to the midpoint of the leaf was designated at position 0, the “midvein”. Each lateral vein was then allocated a position number in reference to position 0.

## Acknowledgements

With thanks to Samuel Leiboff and China Lunde for extensive help and advice, ARS microscopy staff, UC Berkeley greenhouse staff, and the University of Edinburgh Plant Growth Facilities.

## Author Contributions

SH conceived the original project, developed the mapping population and secured funding. AR and AS designed and carried out the experiments. AR and SH wrote the manuscript. AR agrees to serve as the author responsible for contact and ensures communication.

## Materials and Data Availability

All microscopy data is available via the University of Edinburgh DataShare (https://datashare.ed.ac.uk/handle/10283/3987) in The Plant Shape Lab collection. All next generation sequencing data is available via MaizeGDB or the University of Edinburgh DataShare in The Plant Shape Lab collection. Jupyter notebooks for the bioinformatic analyses are available via Github (https://github.com/ThePlantShapeLab/Hojaloca1).

All materials are available on request from the corresponding author. All other data are in the main paper or the supplement.

**Figures 1-7**

**Supplementary Table 1: Custom SSR mapping primers**

**Supplementary Figures 1-3**

**Supplementary Datasets 1-5**

## Supplementary Datasets

**Supplementary Dataset 1: Broad RNAseq differential gene expression comparison between individuals scored as normal or mutant based on seedling phenotype.**

DeSeq2 output of comparisons between samples with seedling phenotypes scored as wild-type or mutant. This data relates to the results shown in Figure 5. All significantly expressed genes between the pools of mutant and wildtype scored samples (padj <0.05) are listed by their gene identifiers (GID) based on the maize B73 RefGenv3 genome, with matched annotations from Maize, and the best BLAST hit for Arabidopsis and rice. Negative log2FoldChange values indicate down-regulation in mutants.

**Supplementary Dataset 2: RNAseq differential gene expression comparison between individuals in cluster 1 versus cluster 2.**

DeSeq2 output of comparisons between samples defined as cluster 1 or 2 based on the hierarchical clustering. This data relates to the results shown in Figure 6. All significantly expressed genes between the cluster 1 and cluster 2 scored samples (padj <0.05) are listed by their gene identifiers (GID) based on the maize B73 RefGenv3 genome, with matched annotations from Maize, and the best BLAST hit for Arabidopsis and rice.

**Supplementary Dataset 3: RNAseq differential gene expression comparison between individuals in cluster 3 versus cluster 2.**

DeSeq2 output of comparisons between samples defined as cluster 3 or 2 based on the hierarchical clustering. This data relates to the results shown in Figure 6. All significantly expressed genes between the cluster 3 and cluster 2 scored samples (padj <0.05) are listed by their gene identifiers (GID) based on the maize B73 RefGenv3 genome, with matched annotations from Maize, and the best BLAST hit for Arabidopsis and rice

**Supplementary Dataset 4: RNAseq differential gene expression comparison between individuals in cluster 4 versus cluster 2.**

DeSeq2 output of comparisons between samples defined as cluster 4 or 2 based on the hierarchical clustering. This data relates to the results shown in Figure 6. All significantly expressed genes between the cluster 4 and cluster 2 scored samples (padj <0.05) are listed by their gene identifiers (GID) based on the maize B73 RefGenv3 genome, with matched annotations from Maize, and the best BLAST hit for Arabidopsis and rice

**Supplementary Dataset 5: Self-organising map information for all expressed genes**

List of all expressed genes (defined as more than 5 counts in more than 3 samples) with their associated gene identifiers (GID) based on the maize B73 RefGenv3 genome, with matched annotations from Maize, and the best BLAST hit for Arabidopsis and rice, and their node defined in the 4×4 SOM analysis shown in Figure 6D.

---

**Supplementary Figure 1.**
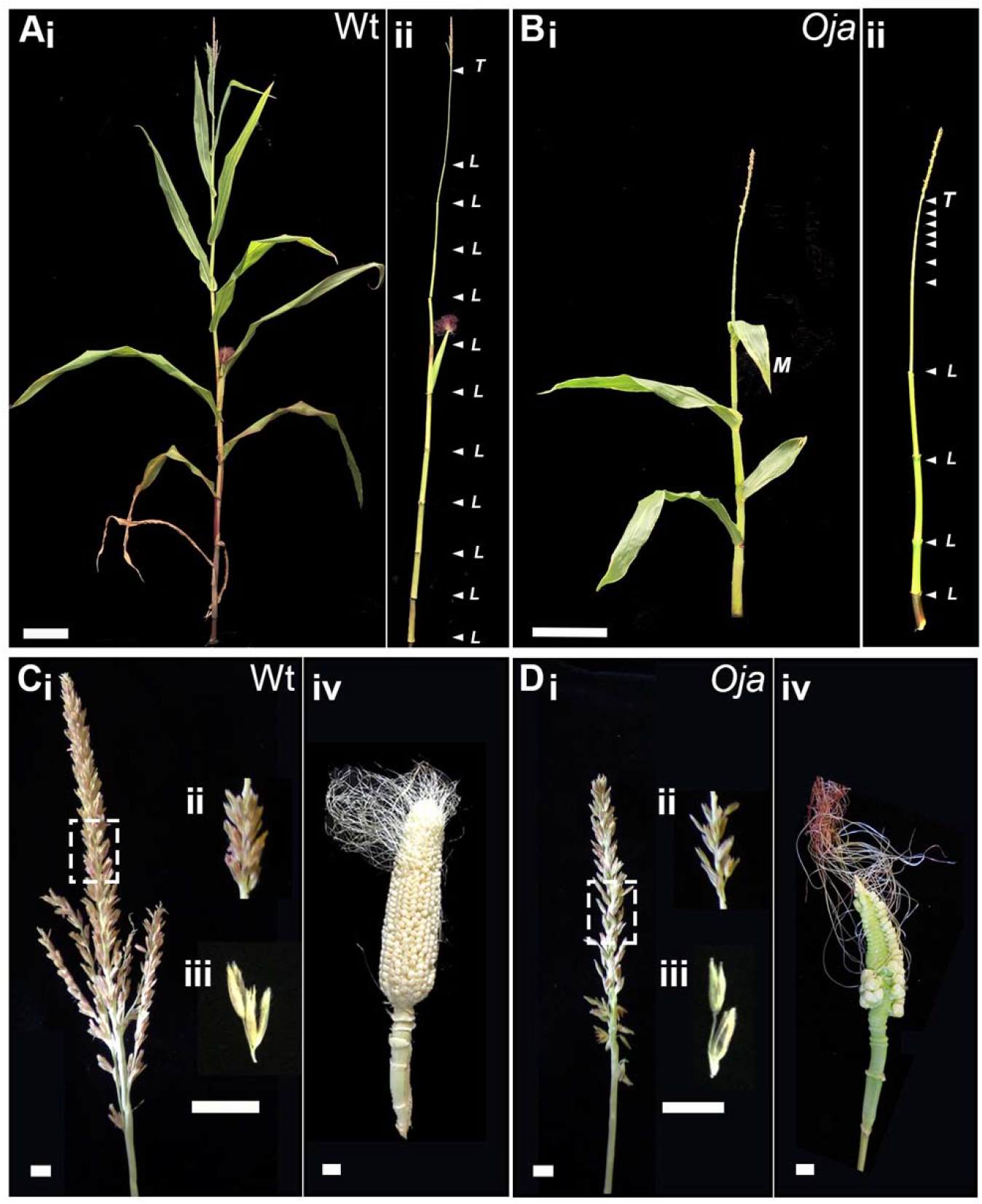
Hoja loca1 Mature Phenotype. The mature phenotype of a wild type (A) and mutant (B) plant, with leaves attached (i) and removed (ii), scale bar is 15cm. The positions of nodes are indicated by the white arrowheads, those that have a leaf are indicated by “L” and the base of the tassel is indicated by “T”. The phenotype of tassels and ears in wild type (C) and mutant (D) plants. (i) Whole tassel, (ii) zoomed-in image of the boxed region, (iii) an individual spikelet pair, (iv) unfertilized ear. Scale bars are 1cm.

**Supplementary Figure 2.**
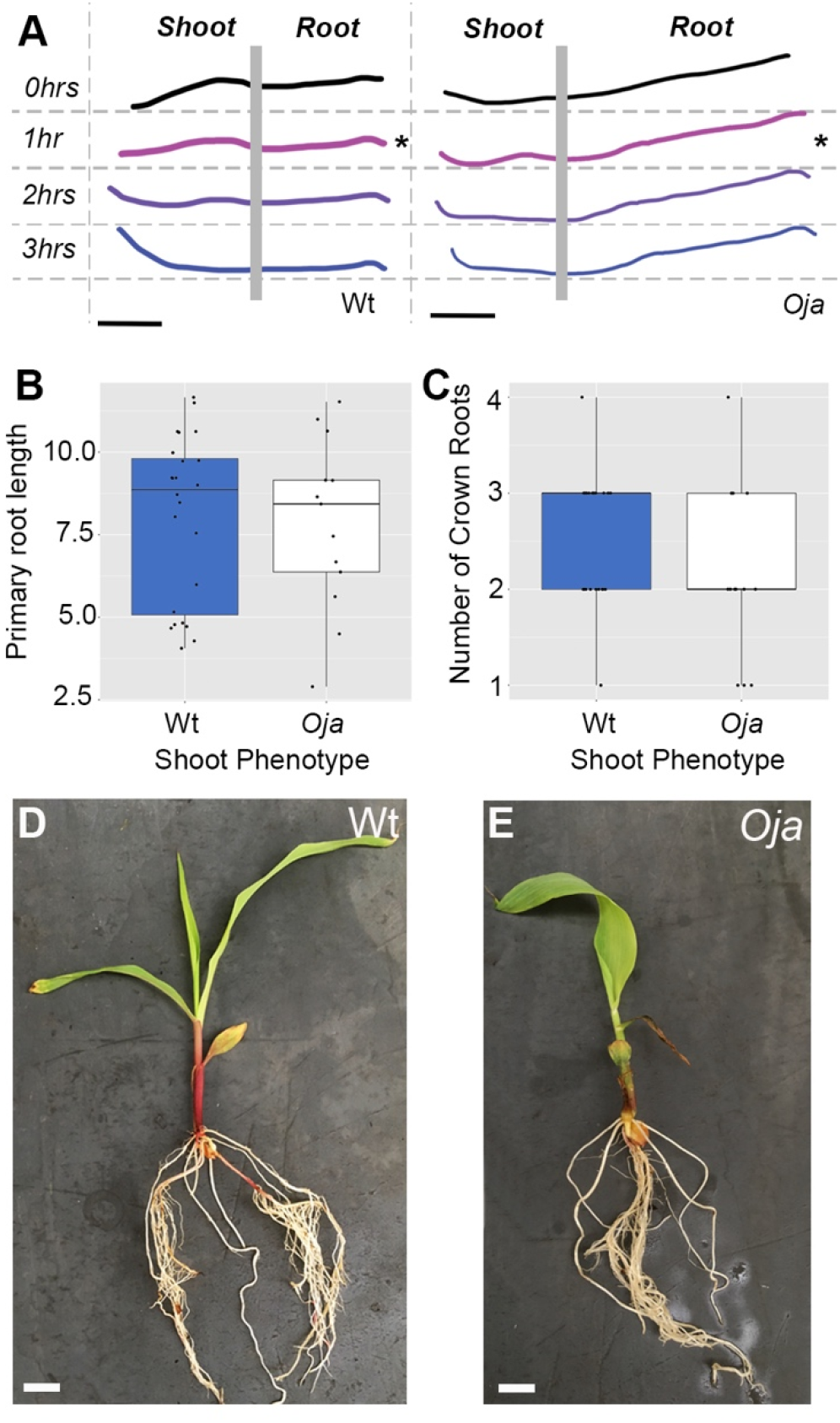
Hoja loca1 mutants have a normal root phenotype. *(A) Oja* seedling roots have a normal gravitropic response, showing a change in root tip orientation by 1 hour after turning (*) (scale bars are 2cm). (B-C) Root architecture in 1.5-week-old seedlings is not significantly different between wild-type and *Oja* plants, as shown by primary root length (B, Welch 2-sample t-test, df=24, n *Oja*=13, Wt=24, p=0.853) and number of initiating crown roots (C, Welch 2-sample t-test, df=19, n *Oja*=13, Wt=24, p=0.1452). (D-E) Root architecture in 3-week-old seedlings appears to be similar between mutant and normal sibling (Wt) (scale bars are 2cm).

**Supplementary Figure 3.**
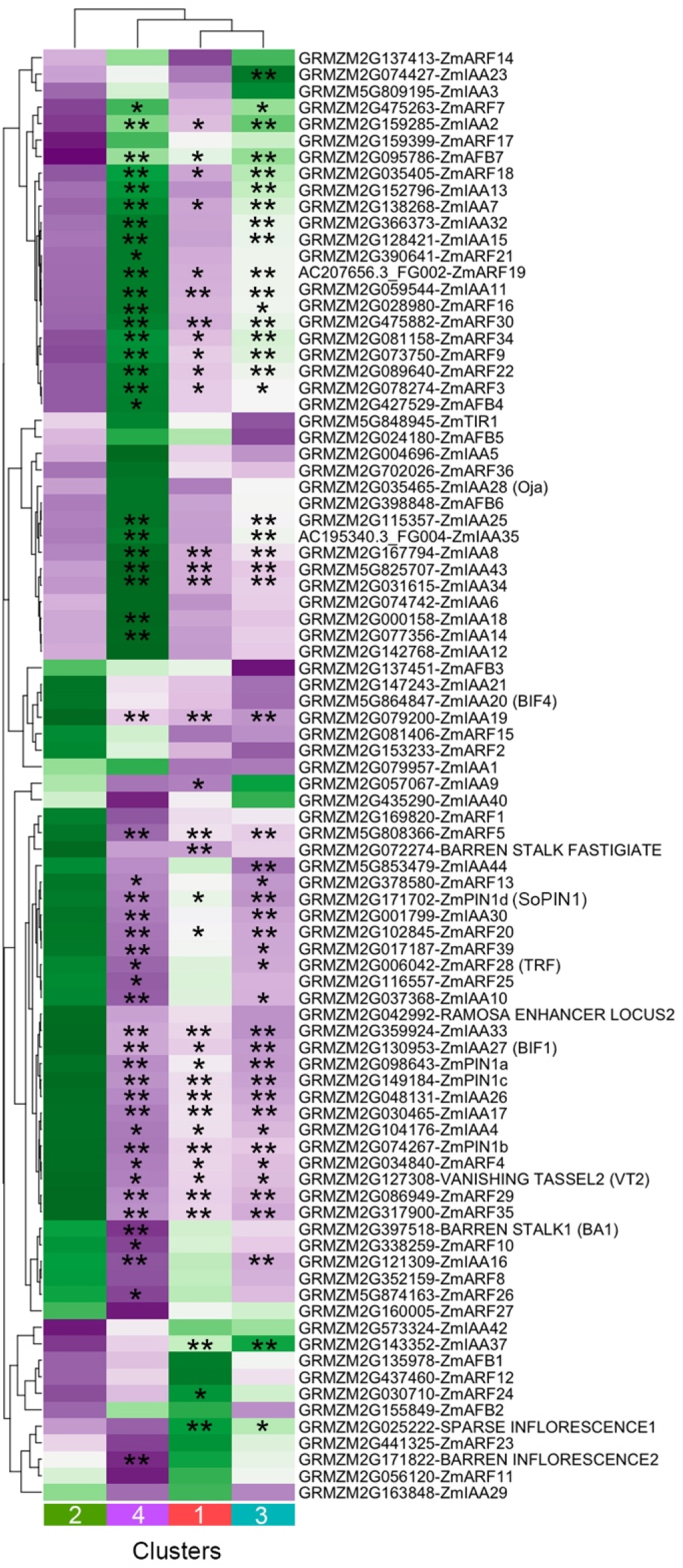
Hoja loca1 mutants show changes in auxin-signaling related genes. Heatmap of average FPKM values of Auxin related genes in clusters 1-4. Colouring is relative average FPKM per row. (* & **) indicate genes that are significantly differentially expressed (padj<0.05), (**) log2foldchange >1 or <-1, foldchange of 2 or 0.5 respectively, (*) log2foldchange <0.59<1 or >-1<0.59, foldchange of 1.5 or 0.5 respectively between the indicated cluster and cluster 2.

**Supplementary Table 1:**
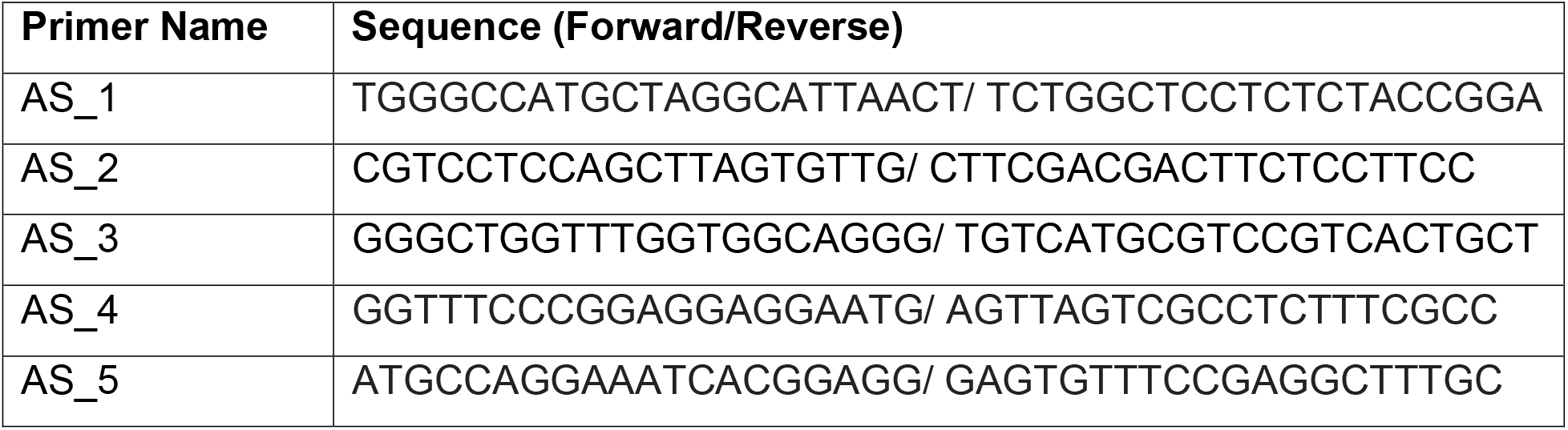
Custom SSR mapping primers. Primers designed using the Gramene SSR prediction tool (Temnykh et al., 2001) to map the *Oja* mutation. Primer locations are indicated in Figure. 4.

